# The *Pseudomonas aeruginosa* complement of lactate dehydrogenases enables use of D- and L-lactate and metabolic crossfeeding

**DOI:** 10.1101/313593

**Authors:** Yu-Cheng Lin, William-Cole Cornell, Alexa Price-Whelan, Lars E.P. Dietrich

## Abstract

*Pseudomonas aeruginosa* is the most common cause of chronic, biofilm-based lung infections in patients with cystic fibrosis (CF). Sputum from patients with CF has been shown to contain oxic and hypoxic subzones as well as millimolar concentrations of lactate. Here, we describe the physiological roles and expression patterns of *P. aeruginosa* lactate dehydrogenases in the contexts of different growth regimes. *P. aeruginosa* produces four enzymes annotated as lactate dehydrogenases, three of which are known to contribute to anaerobic or aerobic metabolism in liquid cultures. These three are LdhA, which reduces pyruvate to D-lactate during anaerobic survival, and LldE and LldD, which oxidize D-lactate and L-lactate, respectively, during aerobic growth. We demonstrate that the fourth enzyme, LldA, performs redundant L-lactate oxidation during growth in aerobic cultures in both a defined MOPS-based medium and synthetic CF sputum medium. However, LldA differs from LldD in that its expression is induced specifically by the L-enantiomer of lactate. We also show that all four enzymes perform functions in colony biofilms that are similar to their functions in liquid cultures. Finally, we provide evidence that the enzymes LdhA and LldE have the potential to support metabolic cross-feeding in biofilms, where LdhA can catalyze the production of D-lactate in the anaerobic zone that is then used as a substrate in the aerobic zone. Together, these observations further our understanding of the metabolic pathways that can contribute to *P. aeruginosa* growth and survival during CF lung infection.

**IMPORTANCE:** Lactate is thought to serve as a carbon and energy source during chronic infections. Sites of bacterial colonization can contain two enantiomers of lactate: the L-form, generally produced by the host, and the D-form, which is usually produced by bacteria including the pulmonary pathogen *Pseudomonas aeruginosa*. Here, we characterize *P. aeruginosa*’s set of four enzymes that it can use to interconvert pyruvate and lactate, the functions of which depend on the availability of oxygen and specific enantiomers of lactate. We also show that anaerobic pyruvate fermentation triggers production of the aerobic D-lactate dehydrogenase in both liquid cultures and biofilms, thereby enabling metabolic cross-feeding of lactate over time and space between subpopulations of cells. These metabolic pathways could contribute to *P. aeruginosa* growth and survival in the lung.

## INTRODUCTION

During growth and survival in communities, bacteria encounter microniches with conditions that differ from those of the external environment. Gradients form over biofilm depth due to consumption of resources by cells closer to the periphery (1–3). Efforts to control biofilm behavior in clinical and industrial settings depend on our understanding of the physiological responses to these unique conditions.

We study the effects of biofilm resource gradients on the physiology of the opportunistic pathogen *Pseudomonas aeruginosa*, a major cause of biofilm-based infections and the most prominent cause of lung infections in patients with the inherited disease cystic fibrosis (4). *P. aeruginosa* can generate ATP for growth via O_2_ and nitrate respiration and, to a limited extent, through arginine fermentation (5, 6). Sufficient ATP to support survival can be generated via (1) pyruvate fermentation (7) or (2) cyclic reduction of endogenously produced antibiotics called phenazines, under conditions in which these compounds are reoxidized outside the cell (8, 9). We have found that some of these pathways for ATP generation also facilitate redox balancing for cells in biofilms and that *P. aeruginosa* modulates its overall community architecture in response to the availability of O_2_, nitrate, and phenazines (1, 10).

Previous work has indicated that phenazine-supported ATP generation and survival in anaerobic cell suspensions depends on reactions associated with pyruvate fermentation and oxidation (7, 9). Furthermore, lactate, a product of pyruvate fermentation, is a major component of cystic fibrosis sputum (11) and a significant carbon and energy source for pathogens and commensals of mammalian hosts (12–15). These observations motivated us to investigate the roles of enzymes that interconvert pyruvate and lactate in *P. aeruginosa* growth and biofilm development, including a previously uncharacterized L-lactate dehydrogenase. We found that this enzyme plays a redundant role in aerobic growth on L-lactate, but that its expression is uniquely and specifically induced by the L-enantiomer of lactate, which is typically produced by plant and mammalian metabolism (16–18). Our studies also show that biofilms grown on pyruvate have the potential to engage in substrate crossfeeding, in which D-lactate produced fermentatively in the anoxic microniche acts as the electron donor for aerobic respiration in the upper, oxic portion of the biofilm. These results help us to further define potential pathways of electron flow in *P. aeruginosa* biofilm cells and the diverse metabolisms that could operate simultaneously in bacterial communities.

## RESULTS AND DISCUSSION

### The *P. aeruginosa* genome encodes four enzymes that interconvert pyruvate and lactate

To initiate our characterization of pathways for pyruvate and lactate utilization in *P. aeruginosa* PA14, we examined the genome for loci encoding lactate dehydrogenases. PA14 contains four genes with this annotation: *ldhA* (PA14_52270), *lldD* (PA14_63090), *lldE* (PA14_63100), and *lldA* (PA14_33860) (**Fig. 1**)., *ldhA* encodes a “lactate dehydrogenase” that reduces pyruvate to lactate during anaerobic pyruvate fermentation (7) (**Fig. 1**). According to computational prediction (19), *ldhA* is co-transcribed with three other genes, including the global regulator *gacS*. *lldD* and *lldE* encode an L-lactate dehydrogenase and a D-lactate dehydrogenase, respectively, and are co-transcribed with *lldP*, which encodes a lactate permease. *lldR*, which encodes a repressor of *lldPDE* expression that is deactivated by either L- or D-lactate, lies adjacent to the *lldPDE* operon and is divergently transcribed (**Fig. 1**) (20). *P. aeruginosa* PA14 encodes a second, uncharacterized L-lactate dehydrogenase that is 44% identical to LldD. Though few other pseudomonad species contain more than one L-lactate dehydrogenase (**Fig. S1B**) (19), the *P. aeruginosa* arrangement of *lldR* and *lldPDE* (**Fig. S1A**) is common within the *Pseudomonas* genus. In the model organism *Escherichia coli*, by contrast, *lldPRD* are arranged in an operon and *lldR* responds specifically to L-lactate (21), while the *E. coli* D-lactate dehydrogenase Dld is encoded separately (**Fig. S1A**) (22).

**FIG 1.**
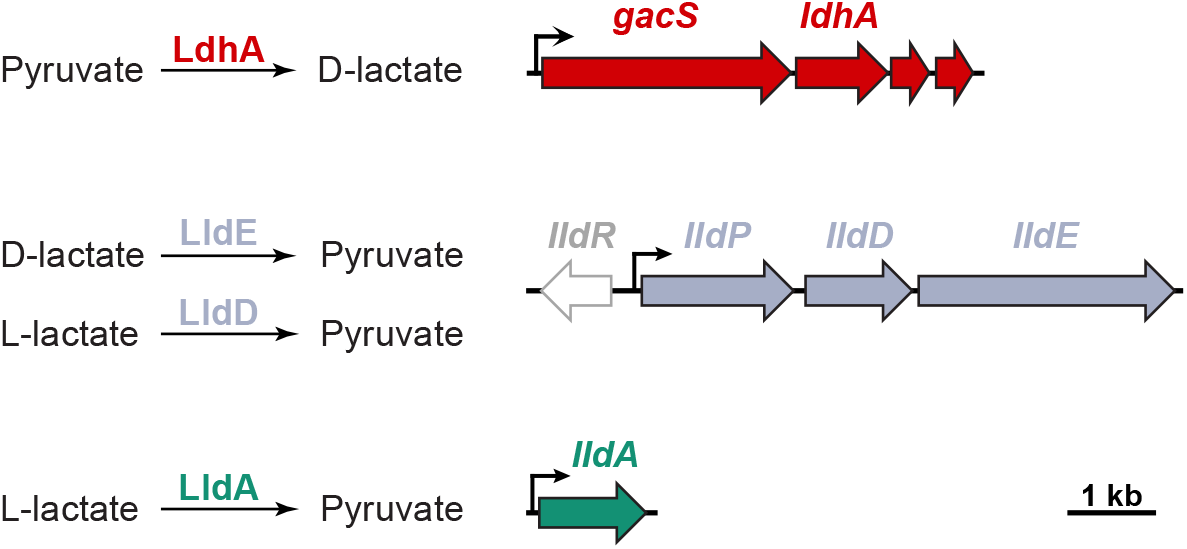
The *P. aeruginosa* genome encodes several enzymes that interconvert pyruvate and lactate. Left: Reactions catalyzed by *P. aeruginosa*’s “lactate dehydrogenases”. Right: Chromosomal loci encoding each of the corresponding enzymes. LdhA catalyzes the reduction of pyruvate during anaerobic survival. LldE catalyzes the oxidation of D-lactate during aerobic growth. Unlike *E coil* which contains only one gene encoding an L-lactate dehydrogenase, *P. aeruginosa* contains two orthologues for this enzyme. LldE catalyzes the oxidation of L-lactate during aerobic growth This study describes a role for LldA in catalyzing the oxidation of L-lactate during aerobic growth

### The *lldPDE* and *lldA* loci are induced by substrates of their respective protein products

Upon noticing the second gene encoding a L-lactate dehydrogenase (i.e., LldA) in the *P. aeruginosa* genome, we wondered whether we could observe its expression under conditions similar to those that induce expression of LldD. We therefore created a suite of *P. aeruginosa* PA14 strains that express GFP under the control of putative promoters for lactate dehydrogenase genes. For the *lldPDE* and *lldA* reporters, we treated ~300 bp of sequence upstream of each of these loci as putative promoters. However, we viewed *ldhA* as a unique case because it was computationally predicted to be co-transcribed with *gacS*. GacS is a notorious, global regulator of quorum sensing (23) and therefore plays a distinct role in *P. aeruginosa* physiology from that of *ldhA.* We examined RNAseq data obtained from PA14 biofilms to identify transcriptional start sites in the *gacS*-*ldhA* region of the genome. This profiling showed a transcriptional start site at ~46 bp upstream of the start codon of *gacS* and steady numbers of reads extending through *ldhA* (**Fig. S2A**), suggesting that *ldhA* expression is driven by the promoter upstream of *gacS*. Nevertheless, we created reporter strains that contained portions of sequence from the regions upstream of either *gacS* or *ldhA*, respectively, so that we could detect any potential, independent expression of *ldhA.*

The reporter strains were grown aerobically in defined media. As would be predicted from the RNAseq profiling, we found that the *gacS* promoter region, but not the putative *ldhA* promoter region, drove expression of the fused *gfp* gene (**Fig. S2B**). Low levels of expression from the *gacS* promoter were observed in aerobic growth on D-glucose, L-lactate, and D-lactate (**Fig. S2B**). A low level of expression was also observed for *lldPDE*, but not *lldA*, on D-glucose (**left panel of Fig. 2**). *lldPDE* expression was strongly induced when either D- or L-lactate was provided as the sole carbon source (**middle and right panels of Fig. 2**). These results are consistent with a previous study in *P. aeruginosa* XMG (20), which showed that LldR responds to both enantiomers of lactate. In contrast, *lldA* expression was not observed on D-glucose or D-lactate and was induced strictly by L-lactate (**Fig. 2**). This finding is consistent with LldA’s predicted physiological function and indicates that it is controlled by a regulator that specifically senses the L-enantiomer.

**FIG 2.**
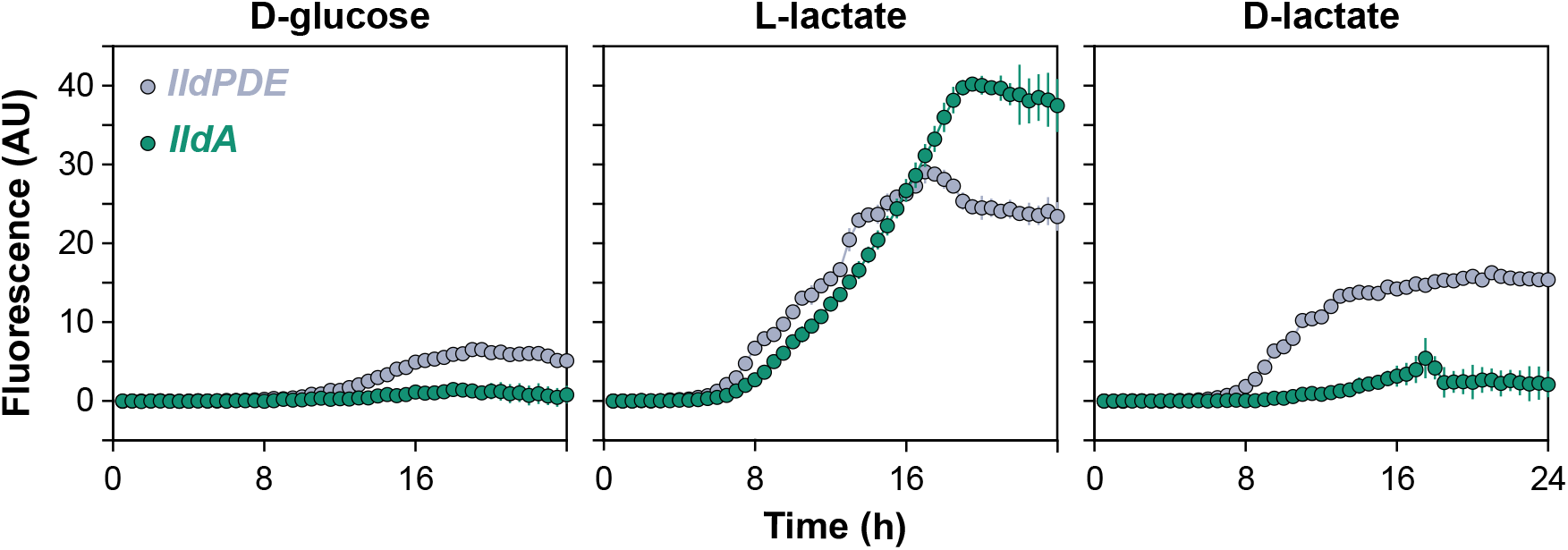
Expression of loci associated with pyruvate and lactate metabolism during aerobic, liquid-culture growth. Strains engineered to express GFP under the control of promoters upstream of *lldP* (which is co-transcribed with *lldD* and *lldE*) or *lldA* were grown in MOPS medium with the indicated compounds provided as sole carbon sources. Error bars, which are obscured by the point marker in most cases, represent the standard deviation of biological triplicates.

### Both LldD and LldA contribute to L-lactate utilization during liquid-culture and biofilm growth

To test whether LldA contributes uniquely to L-lactate utilization, we generated deletion strains lacking *lldDE*, *lldA*, or both loci and tested their abilities to grow aerobically in defined media. When L-lactate was provided as the sole carbon source, deletion of *lldDE* did not completely abolish growth but rather led to biphasic growth at lower rates than that observed for the wild type (**Fig. 3A**). This slow growth of ∆*lldDE* on L-lactate contrasts that seen for an equivalent mutant created in *P. stutzeri* SDM, which shows no growth on L-lactate (24), and indicates that LldD is not the only enzyme that oxidizes L-lactate in *P. aeruginosa*. Indeed, deletion of *lldA* in the ∆*lldDE* background yielded a strain that was completely defective in growth on L-lactate (**Fig. 3A**). However, the Δ*lldA* single deletion did not result in any growth defect on L-lactate (**Fig. 3A**), suggesting that LldA’s activity is redundant with that of LldD. Consistent with findings reported for *P. stutzeri* SDM (24), we found that *lldDE* was necessary for growth with D-lactate (**Fig. 3A**), indicating that LldE is the only enzyme that oxidizes this enantiomer of lactate during aerobic growth.

**FIG 3.**
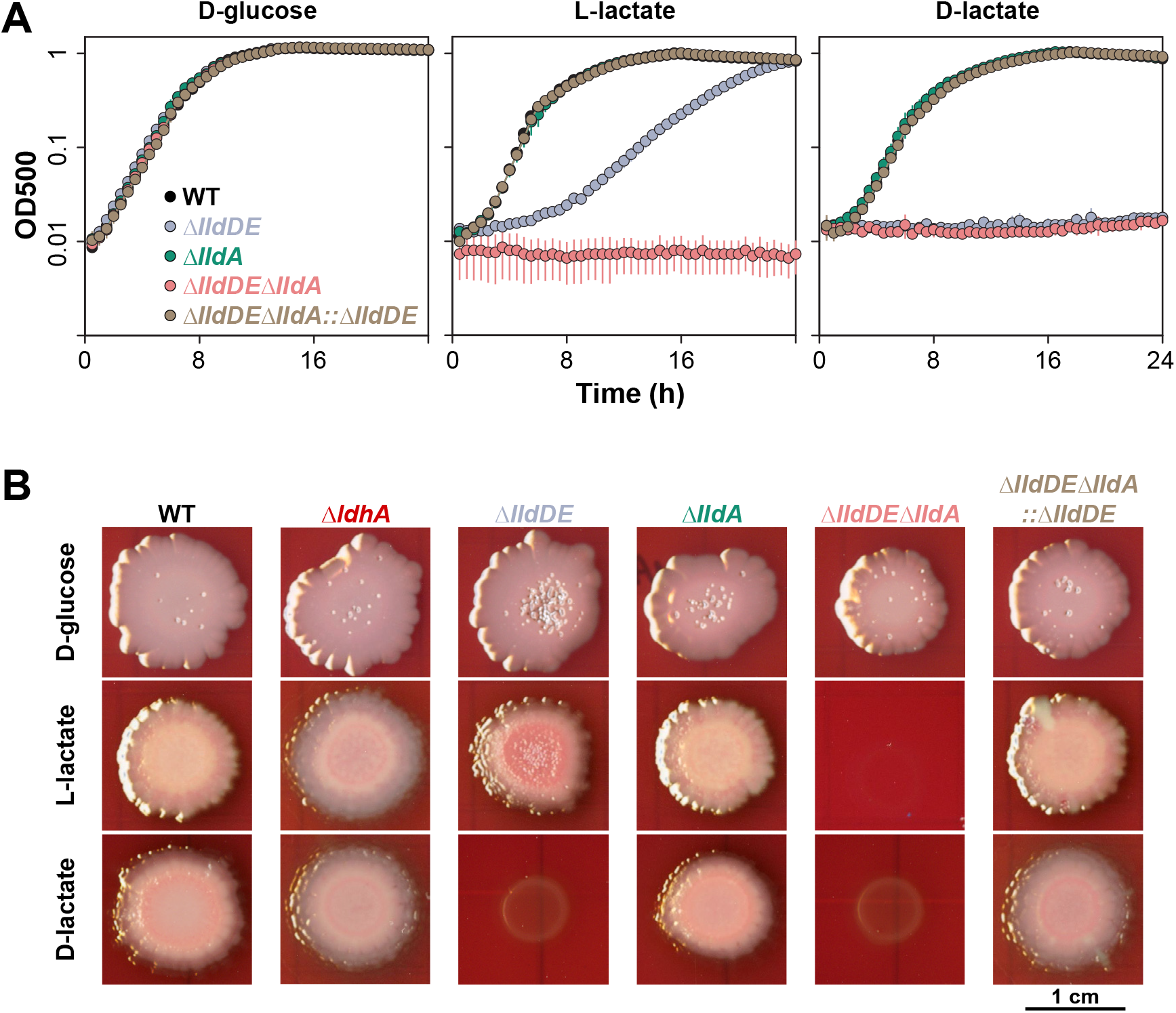
Physiological roles of enzymes that interconvert pyruvate and lactate during growth in shaken liquid cultures and biofilms. **(A)** Aerobic growth of the indicated strains in MOPS medium with D-glucose, L-lactate, or D-lactate provided as the sole carbon source. Error bars, which are obscured by the point marker in most cases, represent the standard deviation of biological triplicates. **(B)** Growth and morphological development of the indicated strains under an oxic atmosphere, on MOPS medium containing the dyes congo red and Coomassie blue and amended with D-glucose, L-lactate, or D-lactate.

The D- and L-enantiomers of lactate are generally associated with bacterial and metazoan metabolism, respectively, with L-lactate being the primary form produced in sites of microbial colonization such as the mammalian gut (13) and the airway of patients with the inherited disease cystic fibrosis (25). We were therefore interested in the relevance of lactate dehydrogenase activity to the growth and morphogenesis of biofilms, which represent a major lifestyle assumed by commensal and pathogenic bacteria. To examine this, we grew deletion mutants lacking lactate dehydrogenase genes as colony biofilms on defined media containing the dyes Congo red and Coomassie blue, which aid visualization of morphogenetic features. Generally, the colony biofilm phenotypes of the mutants matched phenotypes observed during aerobic growth in liquid cultures (**Fig. 3B**). The lack of an effect of *ldhA* deletion (**Fig. 3B**) under these conditions is consistent with the role of LdhA as a pyruvate reductase (7). We note that, in addition to its expected, moderate growth defect on L-lactate, the ∆*lldDE* mutant showed increased production of matrix (as indicated by increased binding of Congo red) (**Fig. 3B**). In other work (1) we have found that this phenotype often correlates with impaired redox balancing and a reduced cellular redox state; in this case, therefore, it may indicate that the LldA enzyme does not sufficiently support maintenance of redox homeostasis in colony biofilms.

### The L-lactate in synthetic cystic fibrosis medium contributes to PA14 growth

Lactate is a major component of cystic fibrosis (CF) sputum (11) and its concentration correlates with exacerbation of chronic infections in CF lungs (25). We predicted that LldD and/or LldA would contribute to PA14 growth in synthetic CF sputum medium (SCFM) (11), which contains 9 mM L-lactate. We grew mutants lacking these L-lactate dehydrogenases in SCFM and found that the double mutant Δ*lldDE*Δ*lldA*, which is not able to grow using L-lactate as a sole carbon source (**Fig. 3**), showed a ~10% decrease in growth (**Fig. 4A**). The growth deficiency was evident after cultures entered late stationary phase (**Fig. 4A** and **Fig. S3**). Both *lldPDE* and *lldA* were transcriptionally induced in SCFM, though *lldPDE* was expressed at a higher level than *lldA* (**Fig. 4B**). These results suggest that LldD and LldA together could contribute to L-lactate utilization for growth in the CF lung environment.

**FIG 4.**
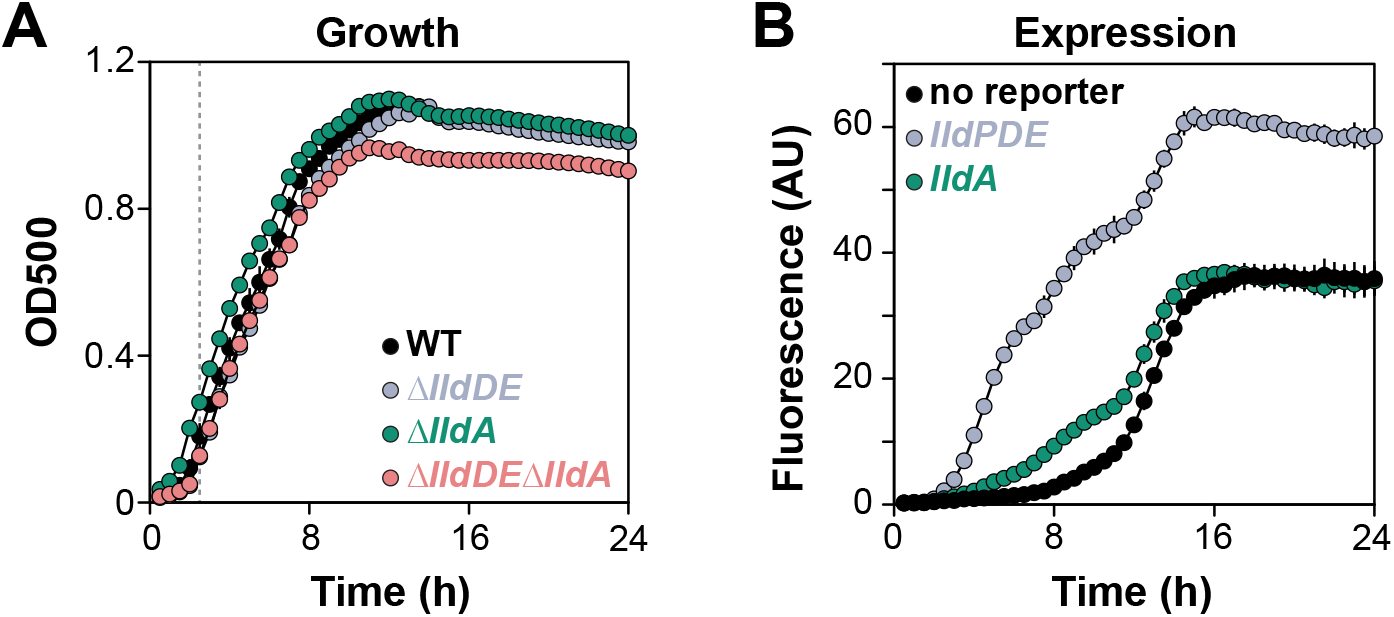
*lldD* and *lldA* utilize the L-lactate in synthetic cystic fibrosis sputum medium (SCFM) for growth. **(A)** Growth of the indicated strains in the SCFM. The vertical dotted line indicates the onset of stationary phase, which is also shown in **Fig. S3**. **(B)** Expression of the indicated reporter constructs in SCFM.

### LldE supports growth of PA14 on self-produced D-lactate

The fact that the *P. aeruginosa* genome encodes enzymes that catalyze the oxygen-dependent interconversion of pyruvate and D-lactate raises the possibility that cells in environments where oxygen availability varies over time or space could engage in metabolic crossfeeding. In this scenario, pyruvate is reduced to D-lactate under anoxic conditions, then D-lactate is oxidized to pyruvate and further metabolized under oxic conditions. We set out to test this possibility in liquid cultures by first incubating *P. aeruginosa* anaerobically with pyruvate (to stimulate D-lactate production), then using the supernatant from these cultures as the medium for aerobic growth of fresh cultures (which should use the D-lactate in an *lldE*-dependent manner) (**Fig. 5A**). First, we verified that *ldhA* supports survival of PA14 under anaerobic conditions in which pyruvate is provided as the major carbon source (**Fig. S4**). We found that this phenomenon required the inclusion of a complex nutrient source (e.g. LB or tryptone) in the medium, possibly as a source of an unidentified cofactor used in pyruvate fermentation. We therefore carried out a series of titration experiments in order to identify a concentration of tryptone that would allow us to observe *lldE*-dependent growth (**Fig. S5A**) and *lldPDE* expression (**Fig. S5B**) at concentrations of lactate that could be produced in the pyruvate fermentation cultures (7). Next, we incubated PA14 cells for 7 days in anoxic, liquid 0.1% tryptone medium containing 40 mM pyruvate, collected cell-free spent medium from these cultures, and used it as the growth medium for freshly inoculated aerobic cultures (**Fig. 5A**). Consistent with the model that PA14- generated D-lactate was present in the medium and acting as a carbon source (**Fig. 5A**), this experimental design yielded high levels of expression from the *lldPDE* reporter (**Fig. 5C**) and revealed a growth defect for the ∆*lldDE* mutant (**Fig. 5B**).

**FIG 5.**
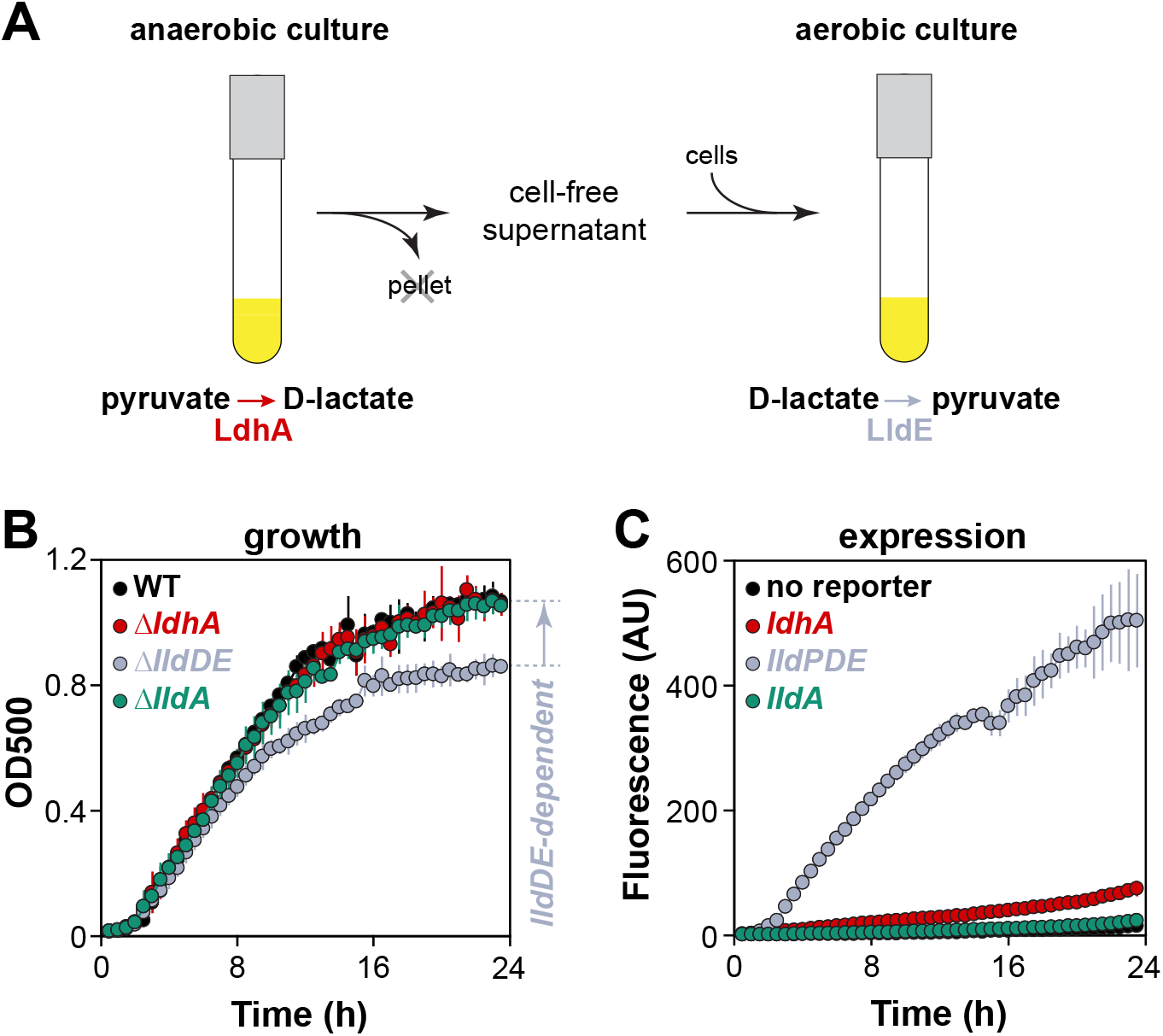
Growth of PA14 on self-produced D-lactate. **(A)** Design schematic for pyruvate/D-lactate crossfeeding experiment in liquid cultures. **(B)** Growth of the indicated strains on supernatant obtained from anaerobic cultures that had fermented pyruvate. The survival medium for the anaerobic cultures was MOPS +0.1% tryptone +40 mM sodium pyruvate. **(C)** Expression of the indicated reporter constructs during growth on the supernatant described in (A).

We next asked whether PA14 has the potential to utilize self-produced D-lactate during growth in colony biofilms. Colony biofilms were first grown aerobically on pyruvate for 2 days to establish sufficient biomass, then transferred to fresh, anoxic plates for 2 days to promote pyruvate fermentation and D-lactate production. These plates were then subsequently moved to oxic conditions and incubated for one day (**Fig. 6A**). This procedure yielded high expression of the *lldPDE* reporter (an L- and D-lactate sensor) but not the *lldA* reporter (an L-lactate-specific sensor) (**Fig. 6B**), indicating the presence of self-produced D-lactate, which could potentially be utilized by PA14 in colony biofilms.

**FIG 6.**
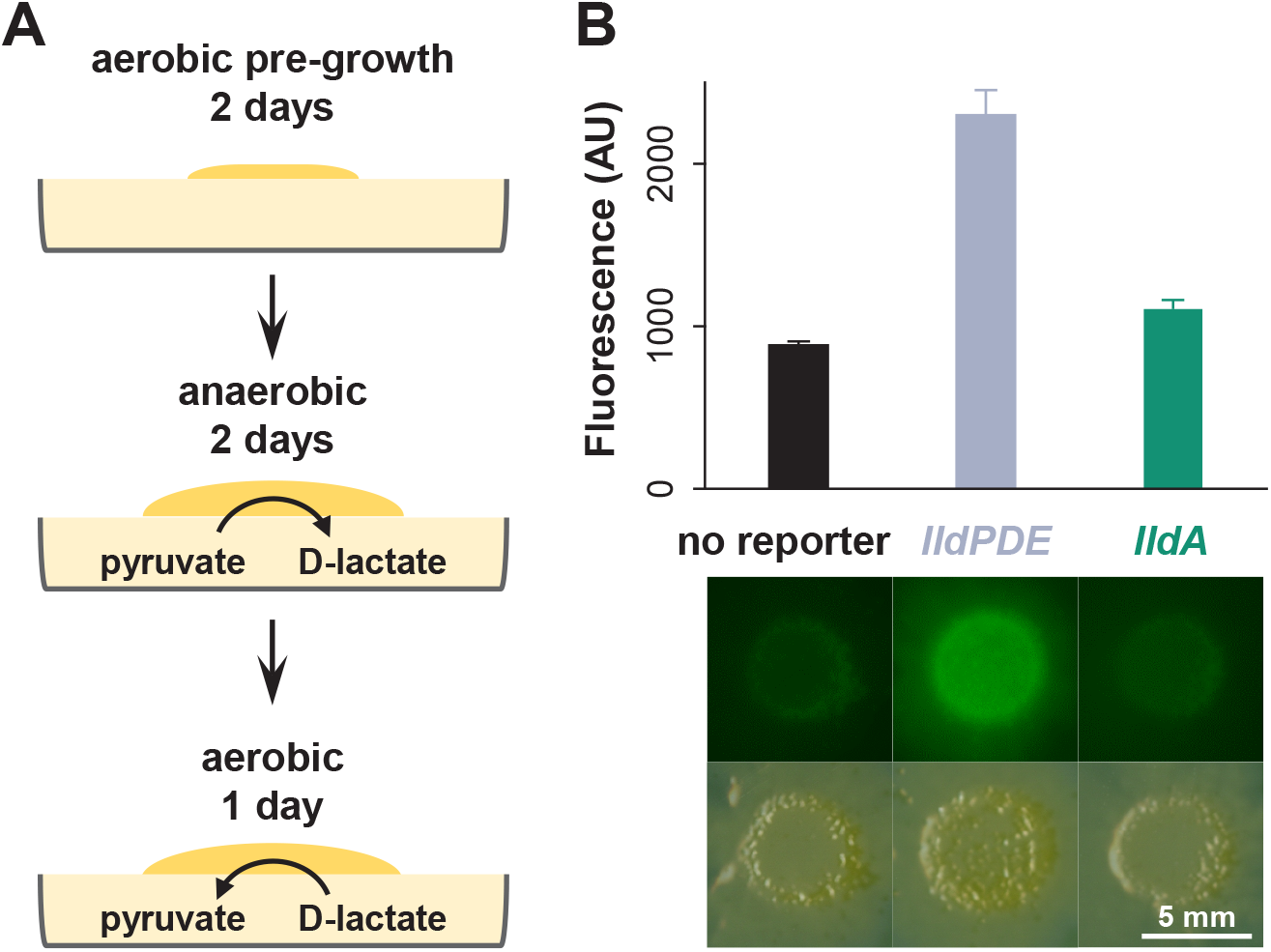
Self-produced D-lactate induces *lldPDE* in biofilms. **(A)** Design schematic for pyruvate/D-lactate crossfeeding experiment in liquid cultures in colony biofilms. Colonies were initially grown atop filter membranes on plates containing 1% tryptone, 1% agar + 40 mM sodium pyruvate for 2 days under an oxic atmosphere to establish biomass, then transferred to fresh plates of medium containing 0.1% tryptone, 1% agar + 40 mM sodium pyruvate in an anoxic chamber to stimulate pyruvate fermentation and D-lactate production for 2 days. Finally, the plates were moved out of the chamber and back into an oxic atmosphere for 1 day. The procedure was carried out at room temperature. **(B)** Fluorescence quantification (top) and images (bottom) of colonies grown using the procedure shown in (A).

### *P. aeruginosa* PA14 cells in biofilms have the potential to engage in pyruvate/lactate crossfeeding over the oxygen gradient

As PA14 colony biofilms increase in thickness, they develop steep oxygen gradients, leading to the formation of metabolic subpopulations in oxic and anoxic zones (26). After observing that PA14 could produce D-lactate under fermentative conditions that acts as an electron donor for aerobic growth, we hypothesized that pyruvate fermentation and aerobic D-lactate utilization could co-occur in biofilms. To test this, we grew colony biofilms on medium with and without pyruvate and examined the expression of the *lldPDE* operon, which is induced by D-lactate. We observed higher expression of *lldPDE* when colonies were grown on medium containing pyruvate (**Fig. 7A**), indicating that cells in the anoxic zone carried out pyruvate fermentation and produced D-lactate. These results suggest that, in addition to generating ATP and supporting survival for cells in oxygen-limited regions of biofilms, pyruvate fermentation can serve to produce D-lactate that supports growth in regions where oxygen is available (**Fig. 7B**). Overall, they add a layer of complexity to our picture of the integrated metabolisms occuring in physiologically heterogeneous bacterial communities.

**FIG 7.**
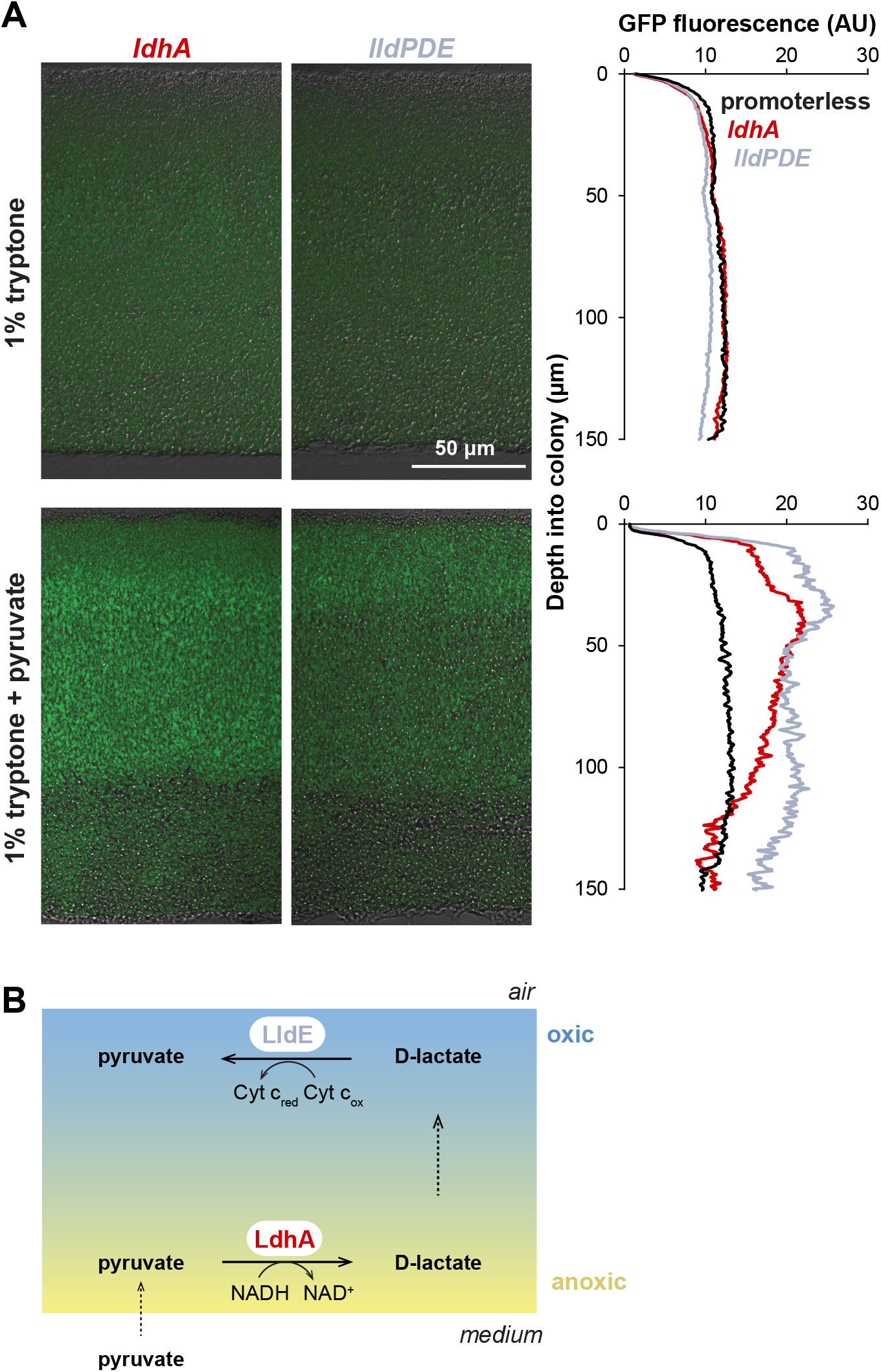
Expression of enzymes for interconversion of pyruvate and lactate over biofilm depth. Expression of *ldhA* (which catalyzes pyruvate reduction under anaerobic conditions) and *lldE*(which catalyzes D-lactate oxidation under aerobic conditions) as reported by promoter *gfp* fusions in thin sections from wild-type PA14 biofilms. Biofilms were grown for 3 days on 1% tryptone medium with or without 10 mM pyruvate. Background fluorescence from a strain containing a promoterless *gfp* is shown at left. Quantification of fluorescence over depth is shown at right. This experiment was repeated in biological triplicate and representative results are shown

### Concluding remarks

Metabolic crossfeeding has been described for polymicrobial communities that are models for those found within the human body and that colonize sites such as the oral cavity and large intestine (27, 28). Our findings indicate that metabolic crossfeeding could take place in a single-species biofilm and arise from physiological heterogeneity, in which a fermentation product formed in the anaerobic microniche is oxidized by cells in the aerobic microniche. Our model for this crossfeeding in biofilms could be viewed as a form of electron shuttling (**Fig. 7B**), with the role of D-lactate as an electron carrier between anoxic and oxic subzones showing some similarities to the that of phenazines (1, 8, 26).

Most pseudomonad species have either *lldD* or *lldA* for L-lactate utilization (**Fig. S1B**). We have shown that, in *P. aeruginosa* PA14, *lldD* and the previously uncharacterized orphan *lldA* encode L-lactate dehydrogenases with redundant functions during growth in liquid cultures. However, *lldA* differs from *lldD* in that it is specifically induced by L-lactate. The presence of redundant L-lactate dehydrogenases in *P. aeruginosa* may represent an adaptation to the host environment, given the fact that host organisms such as humans and plants (18, 29, 30) primarily produce the L-enantiomer. Several studies have implicated L-lactate metabolism in bacterial virulence during infections including *Salmonella*-induced gastroenteritis (13) and gonococcal infection of the genital tract (12). Our results suggest that both LldD and LldA utilize L-lactate in SCFM to promote PA14 growth (**Fig. 4A**). As L-lactate is a significant metabolite available in the CF lung (11, 25, 31, 32), either or both of these enzymes may contribute to *P. aeruginosa*’s ability to colonize and persist in this environment.

## MATERIALS AND METHODS

### Bacterial strains and growth conditions

Unless otherwise indicated, *P. aeruginosa* strain UCBPP-PA14 and mutants thereof (33) were routinely grown in lysogeny broth (LB; 1% tryptone, 1% NaCl, 0.5% yeast extract) (34) at 37°C with shaking at 250 rpm. Overnight cultures were grown for 16 +/-1 hours. For genetic manipulation, strains were typically grown on LB solidified with 1.5% agar. Strains used in this study are listed in **Table S1**. In general, liquid precultures served as inocula for experiments. Overnight precultures for biological replicates were started from separate clonal colonies on streak plates.

### Construction of mutant *P. aeruginosa* strains

For making markerless deletion mutants in *P. aeruginosa* PA14 (**Table S1**) ~1-kb flanking sequences from each side of the target gene were amplified using the primers listed in **Table S2** and inserted into the allelic replacement vector pMQ30 through gap repair cloning in *Saccharomyces cerevisiae* InvSc1 (35). Each plasmid listed in **Table S1** was transformed into *Escherichia coli* strain UQ950, verified by sequencing, and moved into PA14 using biparental conjugation. PA14 single recombinants were selected on LB agar plates containing 100 μg/ml gentamicin. Double recombinants (markerless deletions) were selected on sucrose plates (1% tryptone, 0.5% yeast extract, 10% sucrose, and 1.5% agar). Genotypes of deletion mutants were verified by PCR. Combinatorial mutants were constructed by using single mutants as parent strains.

### Construction of GFP reporter strains

Transcriptional reporter constructs for the genes *ldhA*, *gacS*, *lldP* and *lldA* were made by fusing their promoter sequences with *gfp* using primers listed in **Table S2**. Respective primers were used to amplify promoter regions (as indicated in **Table S1**) and to add an SpeI digest site to the 5’ end of the promoter and an XhoI digest site to its 3’ end. For the *ldhA* reporter, an EcoRI site was used instead of XhoI. Purified PCR products were digested and ligated into the multiple cloning site upstream of the *gfp* sequence of pLD2722, which is a derivative of pYL122 (36) and contains a ribosome-binding site between the MCS and *gfp*. Plasmids were transformed into *E. coli* strain UQ950, verified by sequencing, and moved into PA14 using biparental conjugation. Conjugative transfer of pLD2722 was conducted with the *E. coli strain* S17-1 (36). PA14 single recombinants were selected on M9 minimal medium agar plates (47.8 mM Na_2_HPO_4_, 22 mM KH_2_PO_4_, 8.6 mM NaCl, 18.6 mM NH_4_Cl, 1 mM MgSO_4_, 0.1 mM CaCl_2_, 20 mM sodium citrate dihydrate, 1.5% agar) containing 70 μg/ml gentamicin. The plasmid backbone of pLD2722 was resolved out of PA14 using Flp-FRT recombination by introduction of the pFLP2 plasmid (37) and selected on M9 minimal medium agar plates containing 300 μg/ml carbenicillin and further on sucrose plates (1% tryptone, 0.5% yeast extract, 10% sucrose, 1.5% agar). The presence of *gfp* in the final clones was confirmed by PCR.

### Liquid culture growth assays

Overnight (16-hour) precultures were diluted 1:100 in a clear-bottom, polystyrene black 96-well plate (VWR 82050-756), with each well containing 200 μl of medium. Cultures were then incubated at 37°C with continuous shaking at medium speed in a Biotek Synergy 4 plate reader. Reporter strains were grown in MOPS medium (50 mM MOPS, 43 mM NaCl, 93 mM NH_4_Cl, 2.2 mM KH_2_PO_4_, 1 mM MgSO_4_, 1 μg/ml FeSO_4_ at pH 7.0) amended with one of the following carbon sources: 20 mM D-glucose, 40 mM L-lactate or 40 mM D-lactate (Sigma-Aldrich). Expression of GFP was assessed by taking fluorescence readings at excitation and emission wavelengths of 480 nm and 510 nm, respectively, every 30 minutes for up to 24 hours. Growth was assessed by taking OD readings at 500 nm simultaneously with the fluorescence readings.

### Colony growth assays

Overnight (16-hour) precultures were diluted 1:10 in phosphate buffered saline (PBS). Five μl of diluted cultures were spotted onto MOPS medium (50 mM MOPS, 43 mM NaCl, 93 mM NH_4_Cl, 2.2 mM KH_2_PO_4_, 1 mM MgSO_4_, 1 μg/ml FeSO_4_ at pH 7.0) amended with one of the following carbon sources: 20 mM D-glucose, 40 mM L-lactate or 40 mM D-lactate (Sigma-Aldrich), solidified with 1% agar (Teknova). Colonies were incubated at 25°C for up to 4 days and imaged with an Epson Expression 11000XL scanner. Fluorescence images (**Fig. 6B**) were taken with a Zeiss Axio Zoom.V16 microscope (excitation, 488 nm; emission, 509 nm for imaging of GFP fluorescence).

### Anaerobic survival assays

For each anaerobic liquid sample, 50 μl of overnight (16-hour) precultures were diluted in 5 ml of potassium phosphate (100 mM)-buffered (pH 7.4) LB (7, 38) in a Balch tube and then incubated at 37°C with shaking at 250 rpm for 2.5 h to early-mid exponential phase (OD500 ~0.5). Balch tubes containing subcultures were then transferred into the anaerobic chamber (80% N_2_, 15% CO_2_, and 5% H_2_) and sealed with rubber stoppers to maintain anoxia. Anaerobic tubes were incubated with shaking at 250 rpm at 37°C throughout the period of the survival assay. To measure colony forming units (CFUs) from the anaerobic culture, 10-50 μl of culture were aspirated from the tube with a 1-ml syringe (Care Touch) fitted with a 23G needle (Becton Dickinson), and then serially diluted in LB down to 10^-6^ and plated on 1% tryptone plates for CFU counting.

### Thin sectioning and preparation for microscopic analyses

Thin-sections of *P. aeruginosa* colonies were prepared as described in (39). Briefly, to produce bilayer plates, a 4.5-mm thick bottom layer of medium (1% agar,1% tryptone) was poured and allowed to solidify before pouring a 1.5-mm thick top layer. Precultures were incubated overnight, diluted 1:100 in LB, and incubated until early-mid exponential phase (OD500 ~0.5). Five μl of culture were spotted onto the top agar layer and incubated in the dark at 25°C with >90% humidity (Percival CU-22L) for up to 3 days. Colonies were sacrificed for thin sectioning at specified time points by first covering them with a 1.5 mm-thick layer of 1% agar, which sandwiches each colony between two 1.5 mm layers of solidified agar. Sandwiched colonies were lifted from the bottom layer of agar and soaked in 50 mM L-lysine in PBS (pH 7.4) at 4°C for 4 hours, fixed in 4% formaldehyde, 50 mM L-lysine, PBS (pH 7.4) at 4°C for 4 hours, and then incubated overnight at 37°C. Fixed colonies were washed twice in PBS and dehydrated through a series of ethanol washes (25%, 50%, 70%, 95% ethanol in PBS, 3x 100% ethanol) for 1 hour each and then cleared via three 1-hour washes in Histoclear-II (National Diagnostics HS-202). Cleared colonies were infiltrated with paraffin wax (Electron Microscopy Sciences; Fisher Scientific 50-276-89) at 55°C twice for 2 hours each. Infiltrated colonies were solidified by overnight incubation at 4°C. Sections were cut perpendicularly to the base of the colony in 10-μm slices using an automatic microtome (Thermo Fisher Scientific 905200ER), floated over a water bath at 45°C, collected onto slides, and air-dried overnight. Dried slides were heat-fixed on a hotplate at 45°C for 1 hour, then re-hydrated to PBS by reversing the dehydration steps listed above. Sections were then immediately mounted beneath a coverslip in TRIS-buffered DAPI:Fluorogel (Electron Microscopy Sciences; Fisher Scientific 50-246-93). Differential interference contrast (DIC) and fluorescent confocal images were captured for at least three biological replicates of each strain using an LSM700 confocal microscope (Zeiss).

### RNAseq analysis

Δ*phz* (40) colonies were grown on filter membranes (0.2 μm pore size, 25 mm diameter, Whatman) placed on 1% tryptone, 1.5% agar at 25°C for 76 hours. Colonies were treated with RNAprotect bacteria reagent (Qiagen), samples were excised by microscopic laser dissection, and RNA was extracted using the RNeasy Plant Mini Kit (Qiagen). RNA samples were processed by Genewiz including rRNA depletion and dUTP incorporation for strand-specific sequencing (Illumina HiSeq 2500 platform). Sixteen fastq files were mapped to the reference PA14 genome using Bowtie2 (41) with a ~97% success rate to generate SAM (sequence alignment map) files. SAM files containing ~2 x 10^8^ reads in total were merged, sorted and indexed with SAMtools (42). Read coverage was visualized with Integrative Genomics Viewer (IGV) software (43).

### Preparation of synthetic cystic fibrosis sputum medium (SCFM)

SCFM was prepared as described in (11). SCFM contains the following ingredients: 2.28 mM NH_4_Cl, 14.94 mM KCl, 51.85 mM NaCl, 10 mM MOPS, 1.3 mM NaH_2_PO4, 1.25 mM Na_2_HPO_4_, 0.348 mM KNO_3_, 0.271 mM K_2_SO_4_, 1.754 mM CaCl_2_, 0.606 mM MgCl_2_, 0.0036 mM FeSO_4_, 3 mM D-glucose, 9.3 mM sodium L-lactate, 0.827 mM L-aspartate, 1.072 mM L-threonine, 1.446 mM L-serine, 1.549 mM L-glutamate•HCl, 1.661 mM L-proline, 1.203 mM Glycine, 1.78 mM L-alanine, 0.16 mM L-cysteine•HCl, 1.117 mM L-valine, 0.633 mM L-methionine, 1.12 mM L-isoleucine, 1.609 mM L-leucine, 0.802 mM L-tyrosine, 0.53 mM L-phenylalanine, 0.676 mM L-ornithine•HCl, 2.128 mM L-lysine•HCl, 0.519 mM L-histidine•HCl, 0.013 mM L-tryptophan, 0.306 mM L-arginine•HCl. Depending on the solubility of the various salts, concentrations of their stock solutions ranged from 0.2 M to 1M. Stock concentrations for D-glucose and sodium L-lactate were 1 M, for amino acid stocks 0.1 mM. No stock solution was prepared for L-tyrosine or L-tryptophan due to poor solubility. The pH of SCFM was adjusted to 6.5 with KOH and sterilized by filtration (Thermo Scientific^TM^ Nalgene^TM^ Rapid-Flow^TM^).

## ACKNOWLEDGEMENTS

This work was supported by NIH/NIAID grant R01AI103369 and an NSF CAREER award to L.E.P.D.

